# Reprogramming of 3D chromatin domains by antagonizing the β-catenin/CBP interaction attenuates insulin signaling in pancreatic cancer

**DOI:** 10.1101/2023.11.10.566585

**Authors:** Yufan Zhou, Tian Li, Zhijing He, Lavanya Choppavarapu, Xiaohui Hu, Ruifeng Cao, Gustavo W. Leone, Michael Kahn, Victor X. Jin

## Abstract

The therapeutic potential of targeting the β-catenin/CBP interaction has been demonstrated in a variety of preclinical tumor models with a small molecule inhibitor, ICG-001, characterized as a β-catenin/CBP antagonist. Despite the high binding specificity of ICG-001 for the N-terminus of CBP, this β-catenin/CBP antagonist exhibits pleiotropic effects. Our recent studies found global changes in three-dimensional (3D) chromatin architecture in response to disruption of the β-catenin/CBP interaction in pancreatic cancer cells. However, an understanding of the functional crosstalk between antagonizing the β-catenin/CBP interaction effect changes in 3D chromatin architecture and thereby gene expression and downstream effects remains to be elucidated. Here we perform Hi-C analyses on canonical and patient-derived pancreatic cancer cells before and after the treatment with ICG-001. In addition to global alteration of 3D chromatin domains, we unexpectedly identify insulin signaling genes enriched in the altered chromatin domains. We further demonstrate the chromatin loops associated with insulin signaling genes are significantly weakened after ICG-001 treatment. We finally elicit the deletion of a looping of IRS1, a key insulin signaling gene, significantly impede pancreatic cancer cell growth, indicating that looping-mediated insulin signaling might act as an oncogenic pathway to promote pancreatic cancer progression. Our work shows that targeting aberrant insulin chromatin looping in pancreatic cancer might provide a therapeutic benefit.

## Introduction

Pancreatic cancer is the tenth most common cancer in the United States, with the third highest mortality and the lowest 5-year survival rate, of less than 10% (Siegel et al. 2021). About 90% of pancreatic cancer cases are classified as pancreatic ductal adenocarcinoma (PDAC). Since pancreatic cancer is so lethal, intensive research has been focused on identifying gene signatures to develop new therapeutic approaches. Recent genomic and epigenetic studies have revealed that epigenetic regulation is an emerging mechanism of PDAC progression (Diaferia et al. 2016; Mao et al. 2017; Roe et al, 2017). Our recent studies found global changes in three-dimensional (3D) chromatin architecture, histone acetylation and methylation levels in response to the inhibition of the β-catenin/CREB Binding Protein (CBP) interaction in PDAC cell lines, demonstrating the plasticity of chromosome architecture and broad epigenomic domains to epigenetic regulators (Gerrard et al. 2019a; Gerrard et al. 2019b). However, the functional crosstalk between 3D chromatin architecture upon antagonizing the β-catenin/CBP interaction and critical downstream signaling pathways in pancreatic cancer are largely unknown.

Wnt/β-catenin signaling is a critical developmental signaling pathway and its dysregulation has been shown to promote development and progression in PDAC (Morris et al. 2010; White et al. 2011). Wnt/β-catenin signaling inhibitors have been extensively investigated both pre-clinically and clinically to treat a variety of cancers (Zhang et al. 2020; Jung and Park, 2020). The specific small molecule inhibitor, ICG-001, was initially characterized as a Wnt/β-catenin signaling antagonist via targeting the CBP/β-catenin interaction and thereby indirectly inhibits CBP’s histone acetyltransferase (HAT) activity (Emami et al. 2004). It has demonstrated *in vivo* activity in various preclinical cancer models and a closely related second generation analog PRI-724 has been evaluated in the clinic (Michael et al. 2014; Manegold et al. 2018; Kahn. 2021). This epigenetic modulator has been further shown to alter gene expression in various cancer cells, including pancreatic cancer cells (Emami et al. 2004, Manegold et al. 2018, Gaddis et al. 2015) as well as to affect the 3D genome and epigenome structure (Gerrard et al. 2019a; Gerrard et al. 2019b). However, a broader exploration of the functional link between 3D chromatin architecture and Wnt/β-catenin signaling inhibitors remains to be elucidated.

Aberrant spatial organization of chromatin has been frequently observed in various cancer cells. For example, altered chromatin architecture has been linked to breast cancer development (Barutcu et al. 2015) and drug resistance (Zhou et al. 2019). Cancer-specific chromatin loops have also been associated with androgen receptor activities in prostate cancer (Rhie et al. 2019). The disruption of histone H1 proteins has been found to initiate 3D chromatin remodeling that promotes lymphoma aggressiveness (Yusufova et al. 2021). Inhibition of Notch signaling or the cell cycle has been shown to alter specific 3D interactions in leukemia (Kloetgen et al. 2020). A recent study has demonstrated that chromatin structure, including compartment, contact domain and loop, have been globally reprogramed during pancreatic cancer metastasis (Ren et al. 2021). However, that study did not explore 3D chromatin-mediated downstream signaling pathways as well as their functional relevance in pancreatic cancer. Thus, it is crucial to fully examine the role of 3D chromatin architecture and its downstream signaling in pancreatic cancer.

To explore the functional relevance of 3D chromatin-mediated signaling in pancreatic cancer, in this study, we conducted *in situ* Hi-C in canonical and patient-derived pancreatic cancer cells to examine the global 3D chromatin changes effected upon the ICG-001 treatment. We then further integrated RNA-seq data to identify distinct chromatin looping genes and their associated signaling pathways. Finally, we illustrated the functional attributes of looping-mediated insulin signaling genes in pancreatic cancer progression.

## Results

### Pancreatic cancer cells are inhibited by Wnt signaling inhibitor ICG-001

In the canonical Wnt signaling pathway, β-catenin by binding to the members of the transcription factor T-cell factor (TCF) family, inhibits differentiation via recruitment of the histone acetyltransferase cyclic AMP-response element binding protein-binding protein (CBP) (Teo et al. 2005). The small molecule inhibitor ICG-001, has been shown to disrupt CBP binding with β-catenin/TCF, which subsequently leads to a recruitment of the highly homologous histone acetyltransferase p300 to bind β-catenin/TCF, thereby initiating differentiation, with corresponding metabolic changes (Hu et al. 2021) (**Figure 1A**). We first conducted *in vitro* functional assays with ICG-001 on the two newly established pancreatic cancer cell lines derived from fresh patient-derived tumor xenografts, PATC53 and PATC50 (Kang et al. 2015), the human pancreatic ductal adenocarcinoma cell line, PANC1, and the human pancreatic epithelial cell line, HPNE. We found that the growth of PANC1, PATC53, PATC50 and HPNE were all significantly inhibited by ICG-001 in a dose dependent fashion at 2, 4, 6, and 12 µM, at Day 4 and 6, compared with the DMSO control (**Figures 1B, S1A-B**). Cell migration in these four cell lines were also inhibited by ICG-001 (**Figures 1C, S2A-B**). Importantly, cell apoptosis was significantly increased in the three pancreatic cell lines, PANC1, PATC53 and PATC50 but not in the normal HPNE cells, upon ICG-001 treatment (**Figures 1D**, **S3A-B**). Since Gemcitabine, a nucleoside analogue, is a standard chemotherapeutic drug for treating pancreatic cancer patients, we also tested the efficacy of the combination of ICG-001 and Gemcitabine and found the cell growth for pancreatic cancer cells was significantly impeded (**S4A-C**). Together, our results demonstrate that ICG-001 inhibited pancreatic cancer cell growth and increased apoptosis as has been previously observed (Manegold et al. 2018).

**Figure 1.**
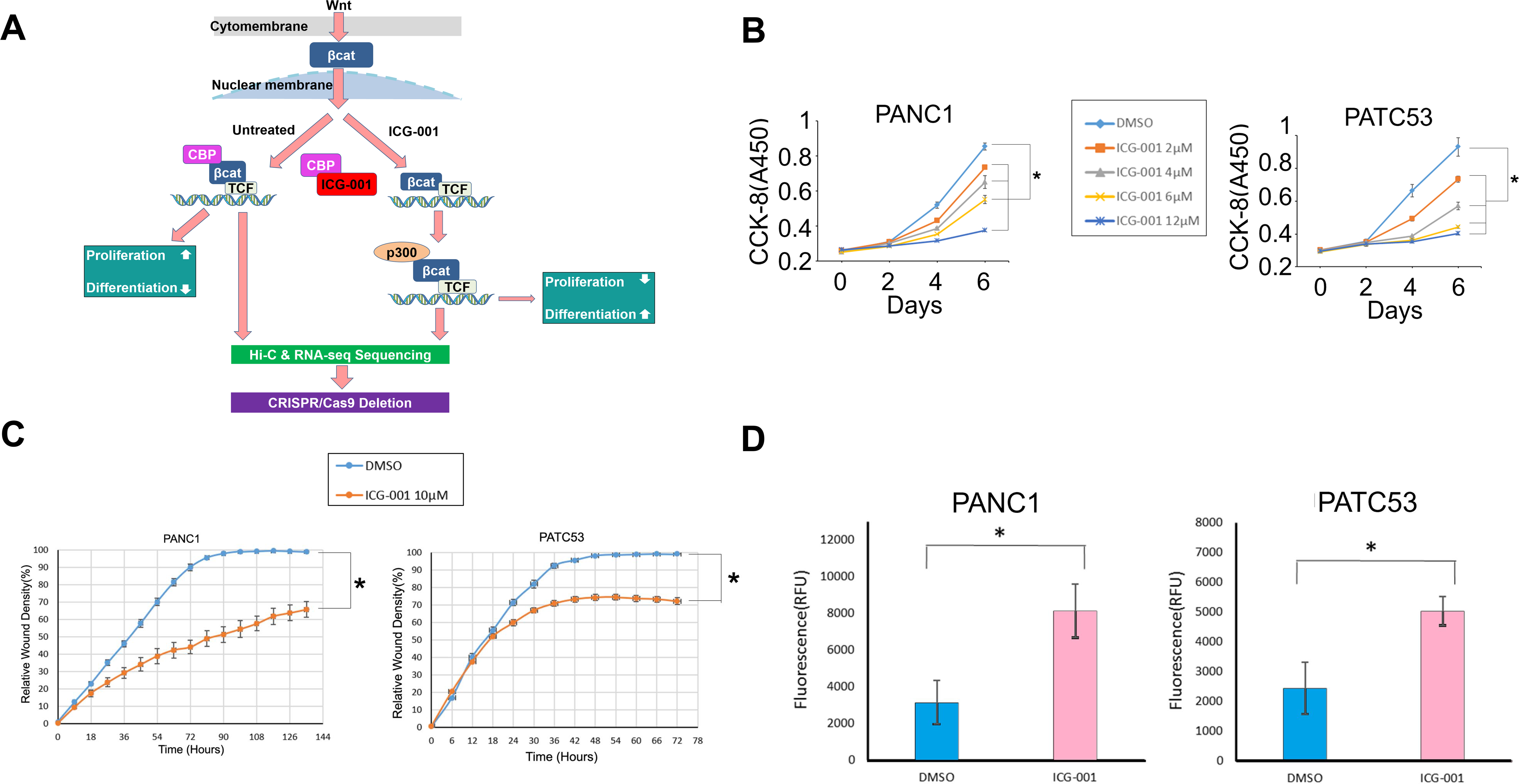
Cell growth curves and apoptosis analysis with the treatment of ICG-001 in pancreatic cancer cells. (**A**) Schematic view of experiments performed in this project. βcat: β-catenin, CBP: histone acetyltransferase cyclic AMP-response element binding protein-binding protein, P300: histone acetyltransferase P300, TCF: T-cell factor. (**B**) Time and dose dependent growth curves of ICG-001 treatment in PANC1 and PATC53. (**C**) Cell migration and invasion assay for PANC1 and PATC53 in the presence of 10 µM ICG-001. (**D**) Apoptosis analysis of pancreatic cancer cell lines PANC1 and PATC53 with ICG-001 treatment. *: Samples t-test, *p* < 0.05.

### The 3D chromatin architecture of pancreatic cancer cells is altered by ICG-001

Previously, we had only investigated the global alteration of 3D chromatin domains by ICG-001 in PANC1 cells (Gerrard et al. 2019a). Therefore, in the present study, we were interested to investigate if ICG-0001’s alteration of 3D chromatin was more broadly affected in pancreatic cancer. To this end, we conducted Hi-C profiling in PATC53 and in PANC1 cells after 48h treatment with DMSO or 10 µM ICG-001 (**Table S1**). We observed clear global compartments changes in PATC53 cells after ICG-001 treatment based on C-score measure (**Figure 2A**) as well as in PANC1 cells. C-score is often used to classify the chromatin compartment type, such that C-score larger or equal to 0 is classified as compartment A, *i.e.*, open, activate chromatin while C-score smaller than 0 is classified as compartment B, *i.e.*, close and repressive chromatin (Zheng et al. 2018). We found that more than half of the compartments flipped after ICG-001 treatment in both pancreatic cancer cell lines (**Figure 2B, Files S1-2**). A bird view of two genomic regions showed two types of compartment change, switched compartment from A to B or B to A and stable compartment from A to A or B to B in PATC53 cells (**Figure 2C**). We then performed the enrichment of KEGG signaling pathway for genes enclosed in the compartment regions with an absolute value of C-score difference larger or equal to 1. Remarkably, the insulin signaling pathway was highly enriched in both PANC1 and PATC53 cells (**Figure 2D**), indicating that it might be a critical downstream signaling regulated by 3D chromatin alteration in response to ICG-001 disruption of the β-catenin/CBP interaction in pancreatic cancer cells.

**Figure 2.**
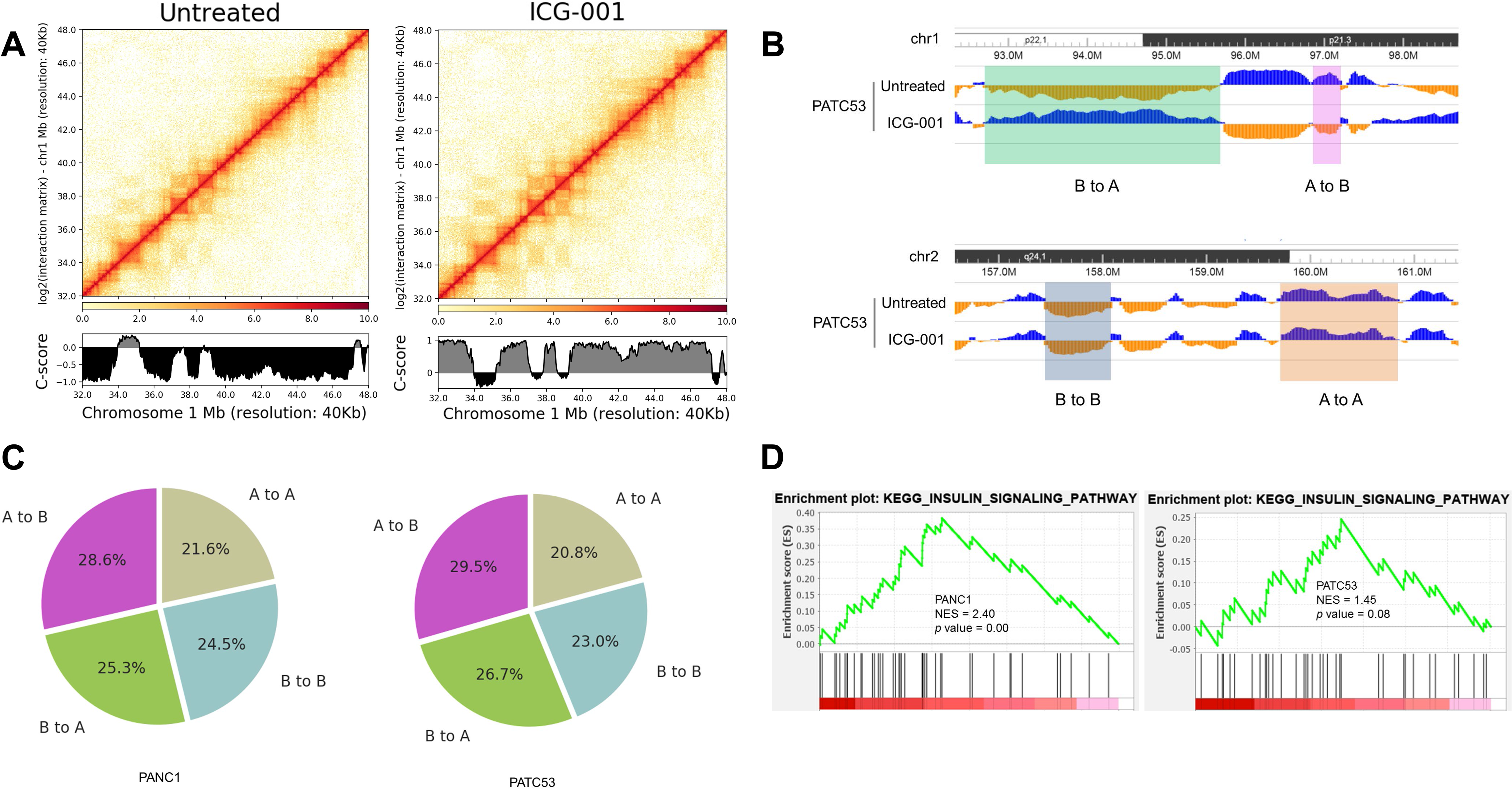
Compartment changes with ICG-001 treatment in pancreatic cancer cells. (**A**) Heatmaps and C-score of Hi-C data in PATC53 cells. (**B**) Illustration of compartment changes. (**C**) The percentage of compartment changes both in PANC1 and PATC53. (**D**) Enrichment of insulin signaling pathway with GSEA from the genes located at regions with absolute value of C-score difference more than 1.

We further examined the Topologically Associating Domains (TADs) before and after ICG-001 treatment and found that the number of and the average sizes of TADs were essentially the same in either treated or untreated conditions (**Figures 3A-B**). However, many of the TADs showed shifted boundaries in both cell lines after ICG-001 treatment (**Figure 3C**). We then conducted RNA-seq analyses (**Table S2**) and identified a total of 5,667 differentially expressed genes (DEGs) in PANC1 or PATC53 cells before and after ICG-001 treatment (**Figures 3D**, **Files S3-4**). Interestingly, some of these DEGs were also shown differentially expressed between PANC1 and HPNE, between PATC53 and HPNE or between PATC50 and HPNE (**Figure 3E**, **Files 5-6**), suggesting some of ICG-001 responsive genes might be directly or indirectly involved in pancreatic tumorigenesis.

**Figure 3.**
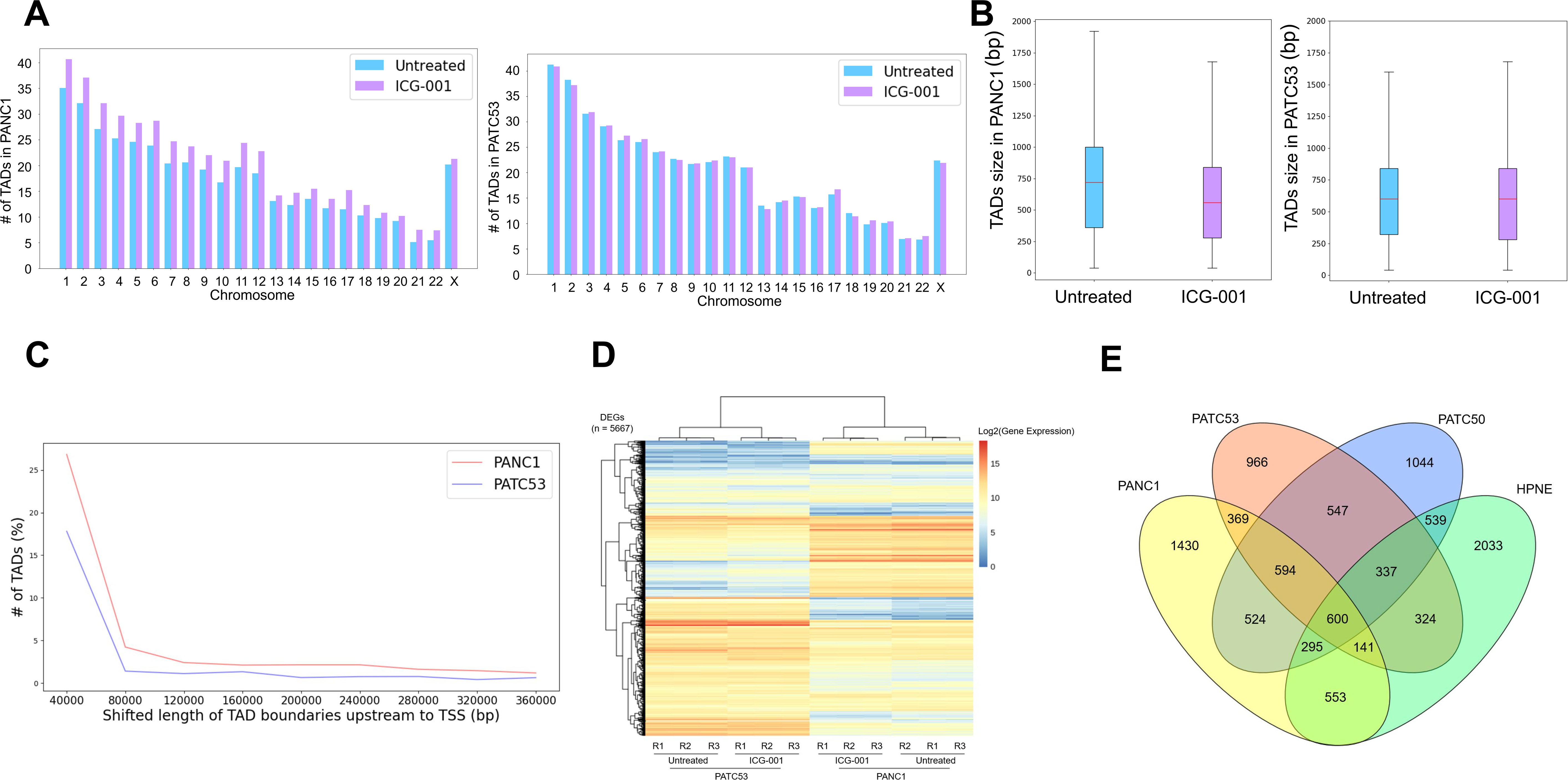
TADs and differentially expressed genes with ICG-001 treatment in pancreatic cancer cells. (**A**) Number of TADs in individual chromosomes in PANC1 and PATC53. (**B**) Size of TADs in PANC1 and PATC53. (**C**) Number of differential TADs in various shifted length of TAD boundaries upstream to TSS after ICG-001 treatment in PANC1 and PATC53. (**D**) Differentially expressed genes (DEGs) with ICG-001 treatment in PANC1 or PATC53 (n = 5667). (**E**) Venn diagram of DEGs in PANC1, PATC53, PATC50 and HPNE.

### Insulin signaling pathway is highly enriched in the altered chromatin structure

Since insulin and insulin receptor substrate 1 (IRS1) are involved in cell proliferation, differentiation and glucose and lipid metabolism, we wanted to further examine the relationship in detail between the insulin signaling pathway (ISP) and 3D chromatin structure (**Figure 4A**). To this end, we examined the difference between the compartments with ISP genes and the compartments without ISP (non-ISP) genes by analyzing the C-score differences. We found that the number of compartment changes were statistically significant between ISP genes and non-ISP genes (**Figure 4B**). ISP genes also showed a higher shifted TAD length than non-ISP genes in both PANC1 and PATC53, respectively (**Figure 4C**). We further investigated the distal-promoter looping events, where the promoter is defined as 5K upstream/downstream Transcription Start Site (TSS) and the distal is defined as either 30-130K upstream of the TSS or 5K downstream of the TSS to Transcription Terminal Site (TTS). We examined the looping changes between before and after ICG-001 treatment as the following four types: (1) ‘None to None’: no looping events in both before and after ICG-001 treatment;(2) ‘Loop to Loop’: having looping events in both before and after ICG-001 treatment; (3) ‘None to Loop’: looping events gained after ICG-001 treatment; and (4) ‘Loop to None’: looping events lost after ICG-001 treatment. For 137 ISP genes, we observed there were 11 ‘None to Loop’ type in PANC1 and 20 in PATC53 and 29 ‘Loop to None’ type in PANC1 and 15 in PATC53, respectively (**Figure 4D**, **File S7-10**). Together, our data illustrated that ISP is highly enriched in the altered chromatin structure.

**Figure 4.**
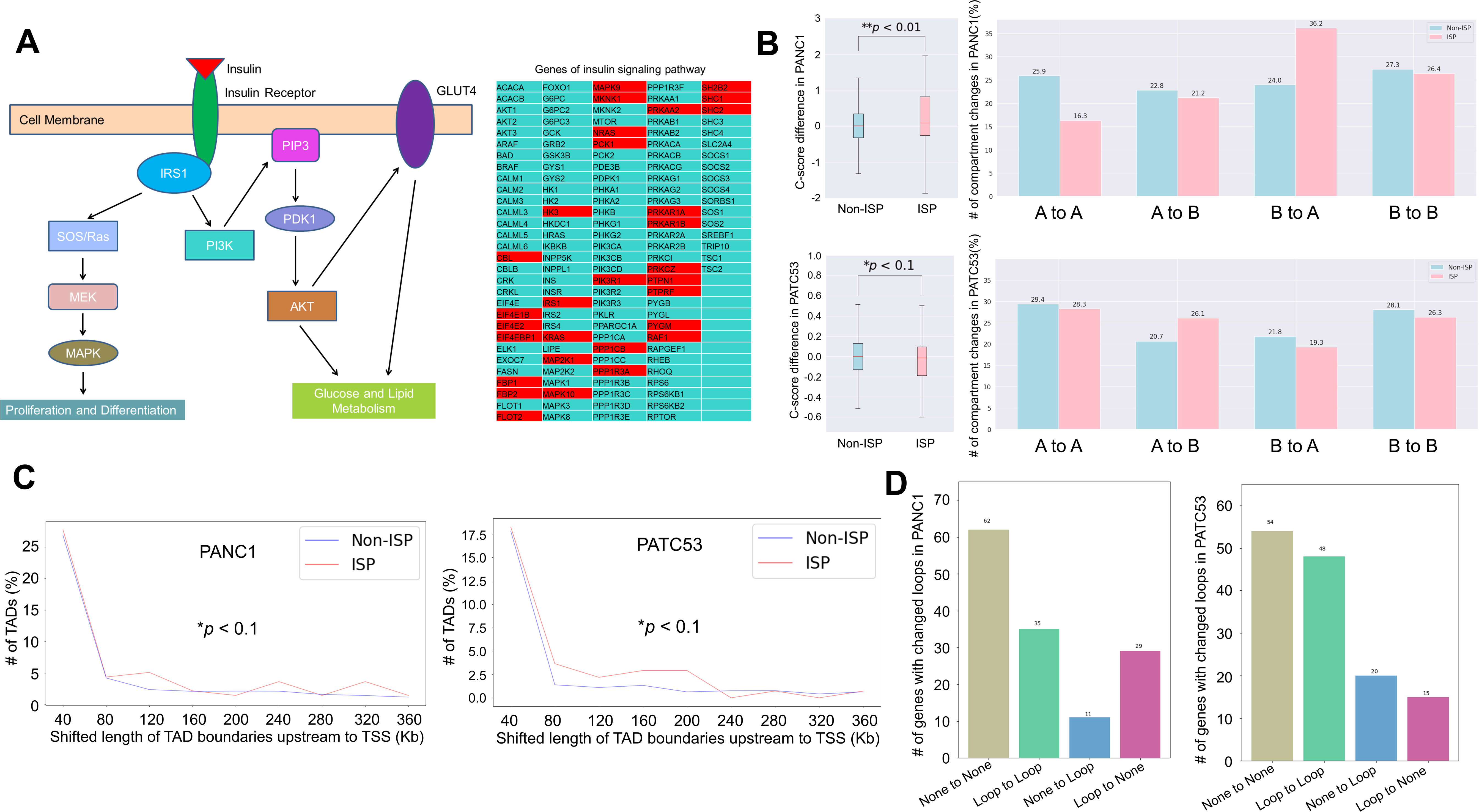
Involvement of insulin signaling pathway in changes of 3D chromatin in pancreatic cancer cells. (**A**) Overview of insulin signaling pathway (ISP). Left: major key molecules in ISP. Right: list of ISP genes, genes with missing looping events after ICG-001 treatment in PANC1/PATC53 are highlighted with red background. (**B**) Comparing of compartment changes of ISP and Non-ISP in PANC1 and PATC53. Left column: C-score difference. Right column: number of compartment changes. * and **: Wilcoxon rank-sum test. ISP: genes of insulin signaling pathway. Non-ISP: genes of not in insulin signaling pathway. (**C**) Number of differential TADs in various shifted length of TAD boundaries upstream to TSS. *: Paired samples t-test. ISP: genes of insulin signaling pathway. Non-ISP: genes of not in insulin signaling pathway. (**D**) Number of genes of insulin signaling pathway with changed loops in PANC1 and PATC53. None to None: no looping events with or without ICG-001 treatment. Loop to Loop: having looping events with or without ICG-001 treatment. None to Loop: looping events gained with ICG-001 treatment. Loop to None: looping events lost with ICG-001 treatment.

### Looping events associated with ISP genes are weakened by ICG-001

We subsequently focused on examining the ISP genes under ‘Loop to None’ type, *i.e.*, the loss of looping events upon ICG-001 treatment. There are a total of 30 common ISP genes in PANC1 or PATC53 showing the loss of looping events (**Figure 5A**). Interestingly, six of these ISP genes were shown to be differentially expressed after ICG-001 treatment, with two up-regulated and four down-regulated (**Figure 5B**). The loss of looping events for these ISP genes was further confirmed by 3C-qPCR experiments in either PANC1 (**Figure 5C**) or PATC53 cells (**Figure 5D**). Remarkably, an experimentally confirmed lost loop for insulin receptor substrate (IRS1), a key ISP gene, is fallen within the shifted TADs in both PANC1 and PATC53 cells (**Figure 5E**), suggesting a functional link between ISP and the alteration of 3D chromatin domains upon the CBP/β-catenin disruption.

**Figure 5.**
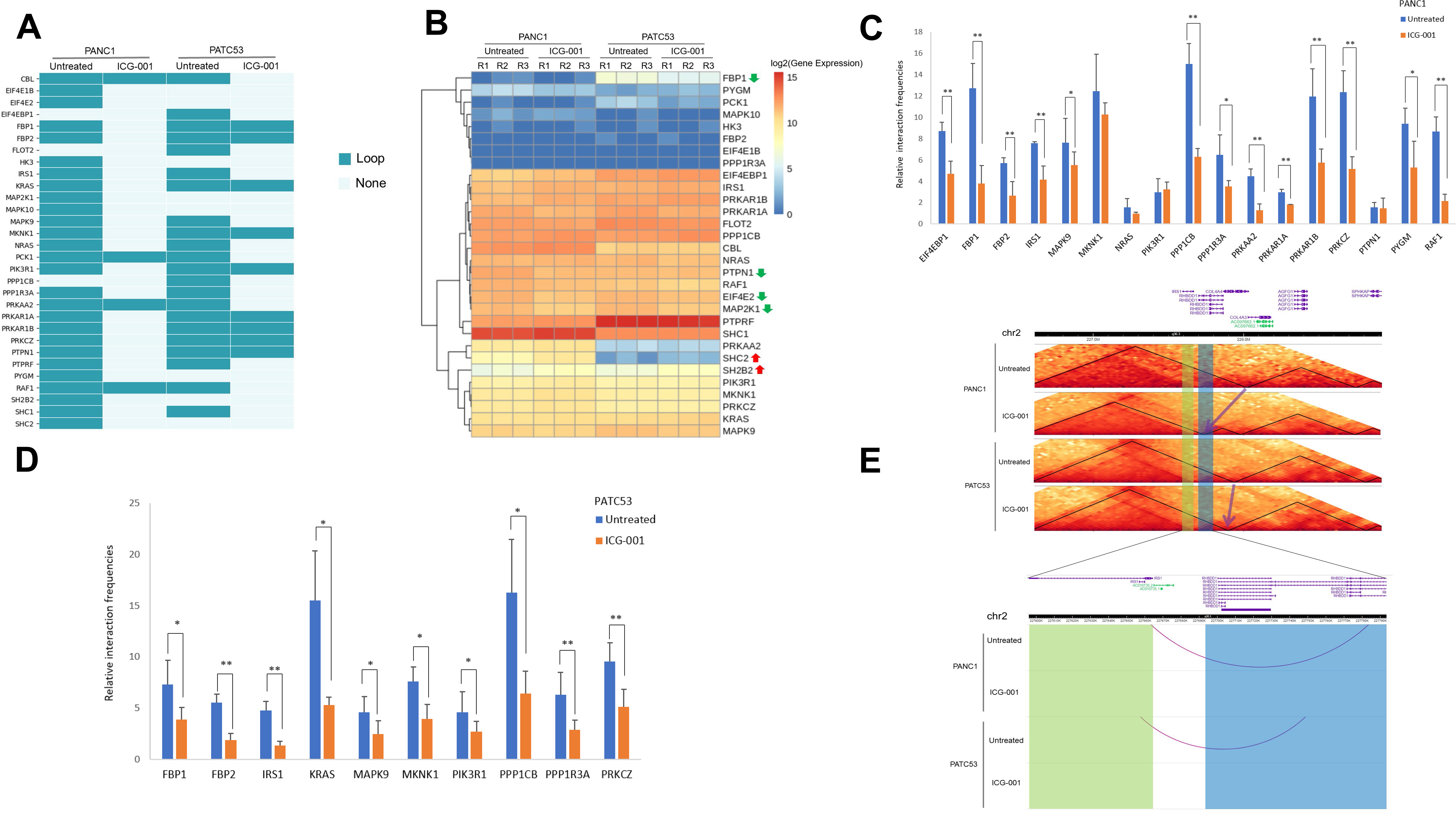
Loss of looping events with ICG-001 treatment in pancreatic cancer cells. (**A**) Loss of looping events of insulin signaling genes in PANC1 or PATC53. Loop: with looping events. None: Without looping events. (**B**) Expression of genes with loss of looping events in insulin signaling pathway in PANC1 and PATC53. Red up arrow: up-regulated genes. Green down arrow: down-regulated genes. (**C**) The relative interaction frequencies of genes of insulin signaling pathway in PANC1 identified by 3C-qPCR experiments. * and **: Samples t-test, *: p < 0.1, **: p < 0.05. (**D**) The relative interaction frequencies of insulin signaling genes in PATC53 identified by 3C-qPCR experiments. * and **: Samples t-test, *: p < 0.1, **: p < 0.05. (**E**) Changes of TADs and looping events with ICG-001 treatment in PANC1 and PATC53. Up panel: TADs changes indicated by arrows. Down panel: looping events indicated by arc. IRS1 gene body is highlighted with yellow. IRS1 upstream 30-130K to TSS is highlighted with blue.

### Deletion of IRS1 distal-promoter looping impedes pancreatic cancer cell growth

To further examine the function of IRS1 distal-promoter looping, we attempted to delete the distal region of 105K length upstream of IRS1 (Chr2: 227,692,583-227,797,349, about 29K far away from TSS of IRS1) by CRISPR/Cas9 technology in PATC53 cells (**Figure 6A**). PCR products showed that the distal region on one allele was successfully deleted in two clones, Del-01 and Del-02 (**Figure 6B**). Sanger sequencing further confirmed such deletion of this genomic distal region (**Figure 6C**). The mRNA expression of IRS1 in both Del-01 and Del-02 clones was also significantly reduced (**Figure 6D**). Furthermore, the protein expression of IRS1 was clearly decreased in the Del-02 clone (**Figures 6E-F**). Cell proliferation was also notably slowed in both the Del-01 and Del-02 clones (**Figure 6G**). However, the ISP proteins, including ERK1/2 (MAPK), phosphate ERK1/2, phosphate AKT and AKT, were not significantly downregulated (**Figure S5A**). This may be due to the fact that the level of IRS1 downregulation was not sufficient to alter the AKT and ERK1/2 pathways. In addition, ICG-001 treatment didn’t have an additive inhibitory effect with Del-01 and Del-02 mutation (**Figure S5B**). Since IRS1 shares the distal region with its neighboring gene RHBDD1, we wanted to rule out that such aberrant pancreatic cancer cell growth was not due to the alteration of RHBDD1. The mRNA levels of RHBDD1 were indeed reduced in both Del-01 and Del-02 clones (**Figure S6A**). The mRNA levels of RHBDD1 were also shown to be downregulated upon the silencing of RHBBD1 (**Figure S6B**). However, the fact that neither cell growth nor IRS1 expression was affected by shRHBDD1 (**Figure S6C-D**) confirmed that RHBDD1 is not involved in IRS1-regulated pancreatic cancer cell phenotypic change. Taken together, our data illustrate that looping-mediated IRS1 expression is able to regulate pancreatic cancer cell phenotypic changes.

**Figure 6.**
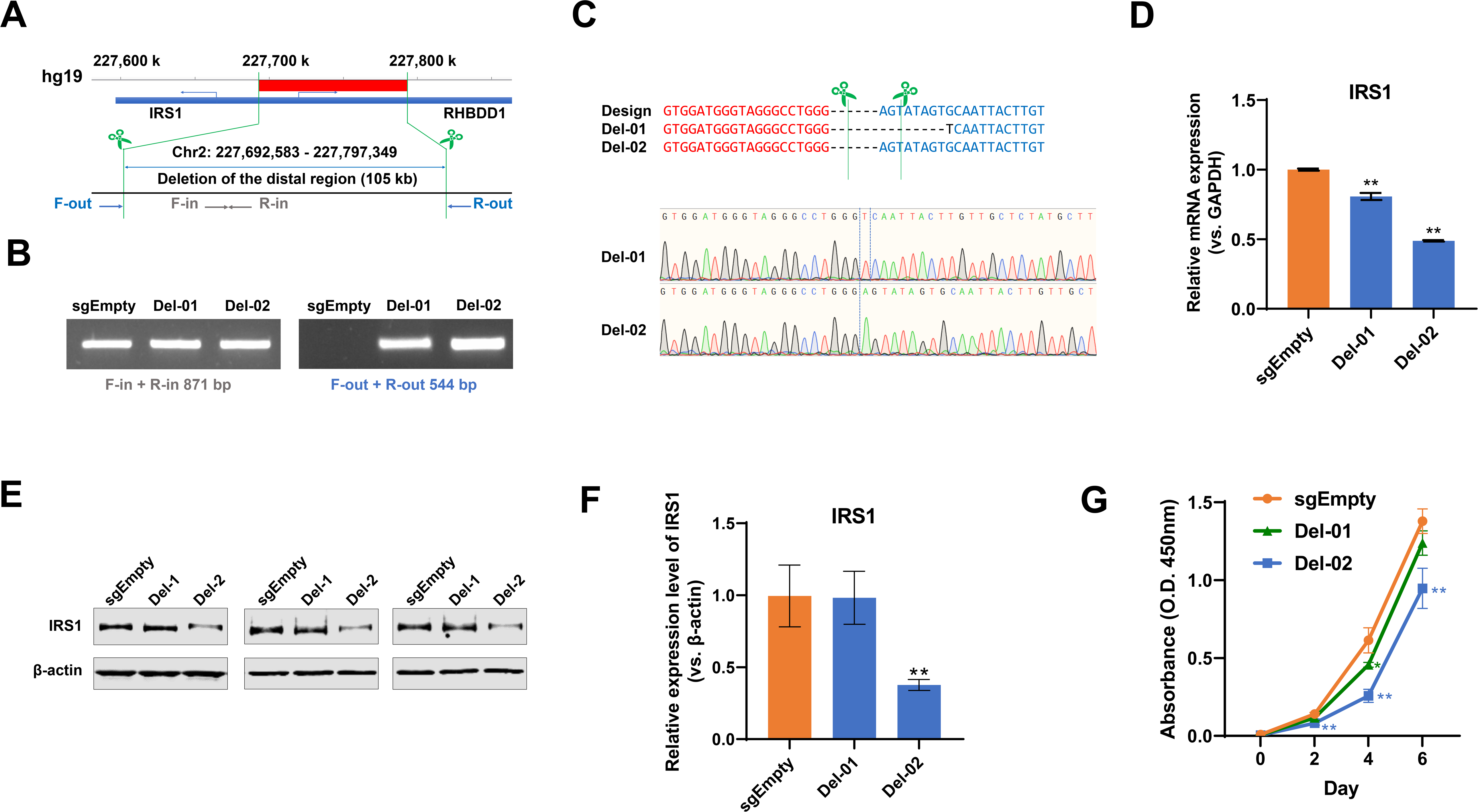
Functional characterization of an IRS1 promoter-enhancer looping in PATC53 cells. (**A**) Design of CRISPR/Cas9-mediated deletion of the enhancer of IRS1. The primer pairs (F/R-out and F/R-in) were used to validate the deletion of enhancer region. (**B**) Gel images of PCR amplification of genomic DNA using primers inside or outside the enhancer region showed the result of the enhancer deletion. Out of 73 clones, 2 clones have shown the validated enhancer deletions. (**C**) The PCR products of the two clones (Del-01 and Del-02) were purified and Sanger sequencing was performed. Sequencing results represented the deletions induced by sgSite-01 and sgSite-02. (**D**) Quantitative PCR was performed to detect the mRNA expression level of IRS1 in enhancer deletion clones. (**E**) Three technical replicates of Western blotting showed the protein expression level of IRS1 in enhancer deletion clones. (**F**) Quantification of protein expression of IRS1 in enhancer deletion clones. (**G**) Cell growth curves showed the growth of Clone Del-02 has significantly decreased comparing with Clone sgEmpty.

## Discussion

Our previous investigations demonstrated that ICG-001, a specific β-catenin/CBP antagonist, was able to sensitize pancreatic cancer tumors to gemcitabine treatment (Emami et al. 2004; Manegold et al. 2018), and to alter chromatin architecture and broad epigenomic domains (Gerrard et al. 2019a; Gerrard et al. 2019b). Another study has also found that ICG-001 altered the expression of DNA replication genes and cell cycle genes such as SKP2 and CDKN1A in pancreatic cancer cells (Arensman et al. 2014). Our current study further systematically examined the effects of disrupting β-catenin/CBP signaling on 3D chromatin architecture in a panel of human pancreatic cancer cells. Interestingly, we identified insulin signaling as the top enriched signaling pathway in the altered chromatin structure, suggesting insulin signaling constitutes a downstream signaling pathway regulated by 3D chromatin looping in response to ICG-001 treatment.

Insulin signaling has been reported to promote the progression of pancreatic cancer (Chan et al. 2014; Deng et al. 2022). Insulin receptor crosstalk with GPCRs stimulates the proliferation of pancreatic cancer (Hao et al. 2017). The insulin-like growth factor receptor IGF1R has been found to contribute to the cancer development (Trajkovic-Arsic et al. 2013), although phase III clinical trials targeting IGF1R inhibition have been unsuccessful (Yee. 2012; Mutgan et al. 2018). One explanation could be that endocrine-exocrine signaling beyond insulin is more essential in the progression of obesity-associated pancreatic cancer (Chung et al. 2020). Our study provides new evidence that the regulation of 3D chromatin structure and its crosstalk with insulin signaling are critical to pancreatic cancer development and progression.

Dysregulated insulin signaling is known to be the primary cause in the development of diabetes mellitus leading to long-term high blood sugar or hyperglycemia, in both type 1 diabetes (T1D) and type 2 diabetes (T2D) (Roloz and Stambolic. 2015). Indeed, both T1D and T2D patients have a two-fold increased risk of developing pancreatic cancer (Stevens et al. 2007; Batabyal et al. 2014). One potential mechanism of hyperglycemia leading to cancer has been reported in which high glucose conditions promote the post-translational modification of O-GlcNAcylation, resulting in nucleotide imbalance, which triggers KRAS mutation in pancreatic cancer (Hu et al. 2019). Pancreatic stellate cells normally have the ability to migrate when they are activated, however, they settle down in the pancreatic islets due to lipotoxicity in T2D (Zhou et al. 2019), thereby boosting the proliferation of pancreatic cancer cells (Liu et al. 2019; Marzoq et al. 2019). Our current findings may provide an opportunity to further examine the role of 3D chromatin regulation in the risk of pancreatic cancer incidence in diabetes patients.

Although looping-mediated IRS1 has been focused in this study, it is worth knowing that the six genes related to insulin signaling (**Figure 5B**) and shown both looping and mRNA expression changes have been reported to be involved in insulin related functions. For example, FBP1 (Fructose-Bisphosphatase 1) has a function in regulating glucose sensing and insulin secretion of pancreatic beta-cells, and PTPN1 (Protein Tyrosine Phosphatase Non-Receptor Type 1) is a negative regulator of insulin signaling by dephosphorylating the phosphotryosine residues of insulin receptor kinase. Therefore, in our future studies, we will examine their looping-mediated functionality in regulating pancreatic cancer progression.

Taken together, we conclude that the disruption of the β-catenin/CBP interaction, thereby indirectly inhibiting CBP’s histone acetyltransferase (HAT) activity with the potential recruitment of p300 HAT activity (Kahn. 2021), attenuates 3D chromatin-mediated insulin signaling in pancreatic cancer cells. Our data indicate that aberrant 3D chromatin-mediated insulin signaling might act as an oncogenic pathway to promote pancreatic cancer progression. Our work provides a rationale that targeting insulin chromatin looping might be a potential therapeutic strategy for treating pancreatic cancer.

## Methods

### Cell lines and reagents

PANC1 (male), PATC53 (male), PATC50 (female) and HPNE (male) cells were cultured in Dulbecco’s modified Eagle’s medium (DMEM) supplemented with 10% fetal bovine serum (FBS), 2 mM L-glutamine and 1% penicillin/streptomycin until 90% confluent. Cells were kept in a cell culture incubator with 37°C and 5% CO_2_ until the cells reached 90% confluence.

### Cell proliferation assays

Cell proliferation was assessed using a Cell Counting Kit-8 (CCK-8, Dojindo Laboratories) following the manufacturer’s instructions. A total of 500 PATC53 cells per well were seeded in a 96-well plate. The absorbance at 450 nm was measured at Day 0, 2, 4 and 6 using BioTek ELx800 Absorbance Microplate Reader. 10 μL of CCK-8 solution was added to each well and the plate was incubated for 60 minutes at 37 °C before the measurement.

### Cell migration and invasion assays

Cell migration and invasion assays were performed on Incucyte ZOOM Live Cell Analysis System. The protocols were executed according to the manufacturer’s user manual about 96-well scratch wound cell migration and invasion assays.

### Cell apoptosis analysis

PANC1, PATC53, PATC50 and HPNE cells were respectively treated with ICG-001 (SELLECKCHEM) at the concentration of 10 µM for 48 hours. Caspase 3/7 activities were detected by Apo-ONE Homogeneous Caspase-3/7 Assay kit (Promega) according to the manufacturer’s protocols.

### *In situ* Hi-C profiling

*In situ* Hi-C experiments were performed according to the previous study (Yang et al. 2020). Briefly, PANC1 and PATC53 cells were respectively treated with ICG-001 (SELLECKCHEM) at the concentration of 10 µM for 48 hours. Cells were crosslinked with 1% formaldehyde then lysed with lysis buffer. The pelleted nuclei were digested with HindIII restriction enzyme overnight. The digested chromatin was further ligated with T4 DNA ligase after filled with biotin labelled dATP. Biotinylated DNA was sheared to a size of 300-500bp and size-selected by AMPure XP beads. Biotinylated DNA was pulled down with Dynabeads MyOne Streptavidin T1 beads and sheared DNA was repaired. Primers including the sequencing index were used to amplify the libraries. Both pancreatic cancer cell lines were conducted in two biological replicates. The prepared *in situ* Hi-C libraries were then sequenced on Illumina HiSeq3000 platform.

### RNA-seq profiling

PANC1, PATC53, PATC50 and HPNE cells were respectively treated with ICG-001 (SELLECKCHEM) at the concentration of 10 µM for 48 hours. Total RNA was extracted from cells with Quick-RNA Miniprep Kit (ZYMO Research) according to manufacturer’s protocols. The RNA-seq libraries were prepared with Ultra II Directional RNA Library Prep Kit for Illumina (NEB) according to the manufacturer’s protocols. All four pancreatic cancer or normal cell types were conducted in three biological replicates. The prepared RNA-seq libraries were then sequenced on Illumina HiSeq3000 platform.

### CRISPR/Cas9-mediated genomic deletion

sgRNAs were cloned into lentiCRISPR v2 plasmid (Addgene, #52961) following the previously published protocol. All sgRNA sequences are listed **in Supplementary Table S3**. The cloned lentiviral CRISPR plasmids were verified by Sanger sequencing and co-transfected with LV-MAX packaging plasmids mix (Thermo Fisher, #A43237) into 293T cells using Lipofectamine 3000 transfection reagent (Thermo Fisher, #L3000015). Culture medium containing lentivirus particles was collected, filtered, and used for lentivirus transfection into PATC53 cells. Empty lentiCRISPR v2 plasmid lacking expression of any sgRNAs was used as the internal control. All plasmids involved in this study are listed in **Supplementary Table S4**.

To delete the distal region of IRS1 gene, two lentiCRISPR plasmids at two sites flanking the target region were co-transfect in PATC53 cells. After 24 hours, the transfected PATC53 cells were selected with 2 μg/mL puromycin for 72 hours and culture to recover for 24 hours. Then the cells were sorted into 96-well plates with 1 cell/well using limited dilution method. The single cells were grown into colonies, then expanded to obtain clonal population for further analysis. Genomic DNA from monoclonal population cells were extracted using PureLink PCR Purification Kit (Thermo Fisher, #K3100-01). Two pairs of primers were used to perform PCR amplification with Q5 High-Fidelity 2X Master Mix (NEB, #M0492S). All validation primers are listed in Supplementary Table 3. The PCR products were used to validate the deletion of target distal region of IRS1 and the sequences were analyzed by Sanger sequencing. Monoclonal populations with validated deleted target distal region of IRS1 were selected for the various functional assays.

### RT-qPCR and 3C-qPCR

Total RNA from cells was extracted using Quick-RNATM Mini Prep (Zymo Research) according to the manufacturer’s instructions. Quantitative Real-time PCR was performed using a Light Cycler 480 Instrument II real-time PCR system (Roche). The relative expression levels of target genes were determined by the 2^-ΔΔCt^ method using GAPDH as an internal control. The primers are listed in **Supplementary Table S3**. 3C-qPCR experiments were performed as previously described (Zhou et al. 2020). The primers of 3C-qPCR are listed in **Supplementary Table S5**.

### Western blotting

Cells were lysed in RIPA lysis buffer supplemented with 1X Halt protease and phosphatase inhibitor cocktail and 5 mM EDTA (Thermo Fisher Scientific). Protein concentrations were quantified with the Pierce BCA protein assay kit. 30 μg of protein extracts per well were loaded and separated by SDS-PAGE gels. Blotting was performed using a nitrocellulose membrane (LI-COR). After 1 hour blocking with 5% non-fat milk in TBS buffer, the membrane was incubated with primary antibody overnight at 4°C. After washes with TBST, the membrane was incubated with IRDye 680RD Goat anti-Mouse IgG secondary antibody or 800CW Goat anti-Rabbit IgG secondary antibody (LI-COR). Signals were detected Odyssey Imaging Systems (LI-COR) following the manufacturer’s instruction. All antibodies used in Western blotting are listed in **Supplementary Table S6**.

### shRNA plasmid mediated inhibition of RHBDD1 expression

RHBDD1 and control shRNA plasmids were purchased from Santa Cruz Biotechnology (sc-94654-SH, sc-108060). shRNA plasmids were transfected into PATC53 cells following the manufacturer’s protocol. At 24 hours after the transfection, 2 μg/mL puromycin were used to select the stably transfected cells for 72 hours. The RHBDD1 expression of the stably transfected PACT53 cells were quantified by RT-qPCR.

### Hi-C data analysis

Raw Hi-C reads were mapped to human HG19 genome by Bowtie 2 (Langmead and Salzberg. 2012). Mapped paired reads were then filtered and normalized by HiC-Pro (Servant et al. 2015). The compartments were identified by CscoreTool v1.1 to compute C-score at 40K resolution (Zheng et al. 2018). The C-score was filtered with bias > 0.2 and < 5. TADs were identified by matrix2insulation.pl to produce insulation score. The significant chromatin interaction loops were identified by HiSIF (Zhou et al. 2020).

### RNA-seq data analysis

Raw RNA-seq reads were mapped to human HG19 genome by HISAT2 (Kim et al. 2019). The expression counts were computed by htseq-count (Anders et al. 2015). The differentially expressed genes were identified by DESeq2 with cut-off of absolute of log2 fold change > 0.6 and *p* value < 0.01 (Love et al. 2014).

### Enrichment of signaling pathway

Enrichment of signaling pathways was performed with GSEA v4.1.0 on GSEAPreranked module using KEGG gene set database before genes that were ranked by C-score differences (Subramanian et al. 2005).

## Data availability

Except raw Hi-C data for untreated PANC1 were downloaded from ENCODE project with GEO accession number GSE105566 (ENCODE Project Consortium. 2012), raw and processed *in situ* Hi-C data for DMSO or 10 µM ICG-001 two-day treated PANC1 and PATC53 cells are deposited in GEO under accession number GSE197293. Raw and processed RNA-seq data for untreated or 10 µM ICG-001 two-day treated PANC1, PATC53, PATC50, and HPNE cells are deposited in GEO under accession number GSE198444.

## Author contributions

VXJ conceived the project. YZ, TL and ZH conducted the experiments, YZ performed the data analysis. LC assisted in conducting the experiments. VXJ and YZ wrote the manuscript, with all authors including MK, GWS, RC and XH contributing to writing and providing the feedback.

## Acknowledgments

We thank the UTHSA Next Generation Sequencing Facilities for services rendered for production of the Hi-C and RNA-seq data. We would also like to thank J. Fleming at Moffitt Cancer Center for sharing us with their human pancreatic cancer cell lines, PATC53 and PATC50. This project was partially supported by grants from NIH R01GM114142 and U54CA217297.

