## Supplementary material for "Reprogramming of 3D chromatin domains by antagonizing the β-catenin/CBP interaction attenuates insulin signaling in pancreatic cancer": Suppl. Materials

**Supplementary Figure S1. Cell growth curve with the treatment of ICG-001.**

(**A**) Growth curve of PATC50 cells in various concentrations of ICG-001. (**B**) Growth curve of HPNE cells in various concentrations of ICG-001.

**A**

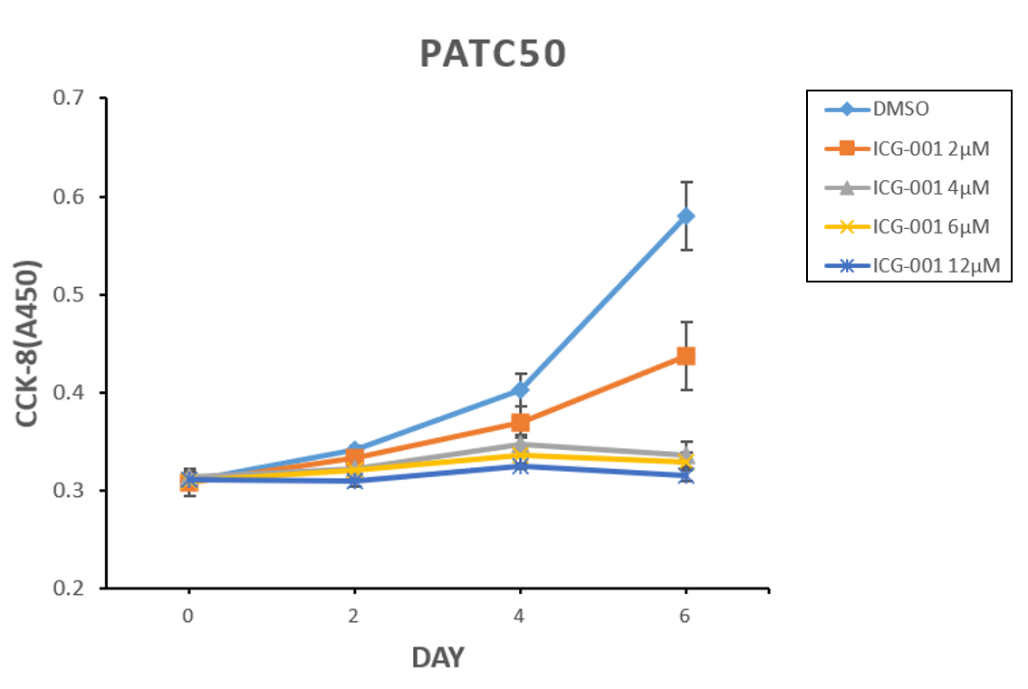

**B**

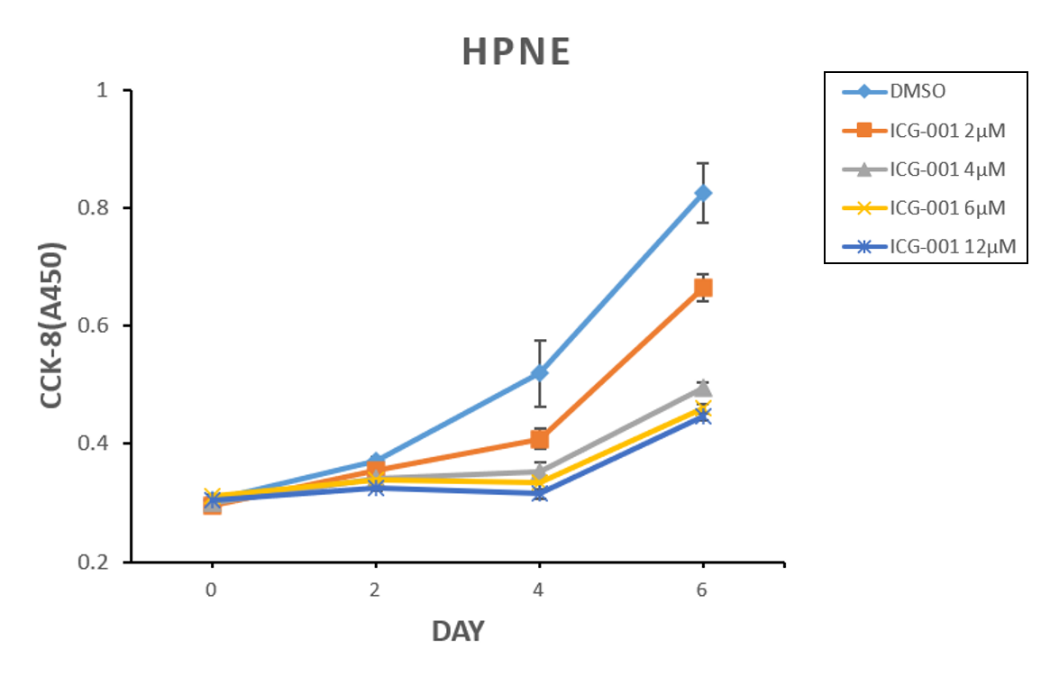

**Supplementary Figure S2. Cell migration and invasion assays upon ICG-001 treatment.**

(**A**) Cell migration and invasion assay for PATC50 in the presence of 10 µM ICG-001. (**B**) Cell migration and invasion assay for HPNE in the presence of 10 µM ICG-001.

**A**

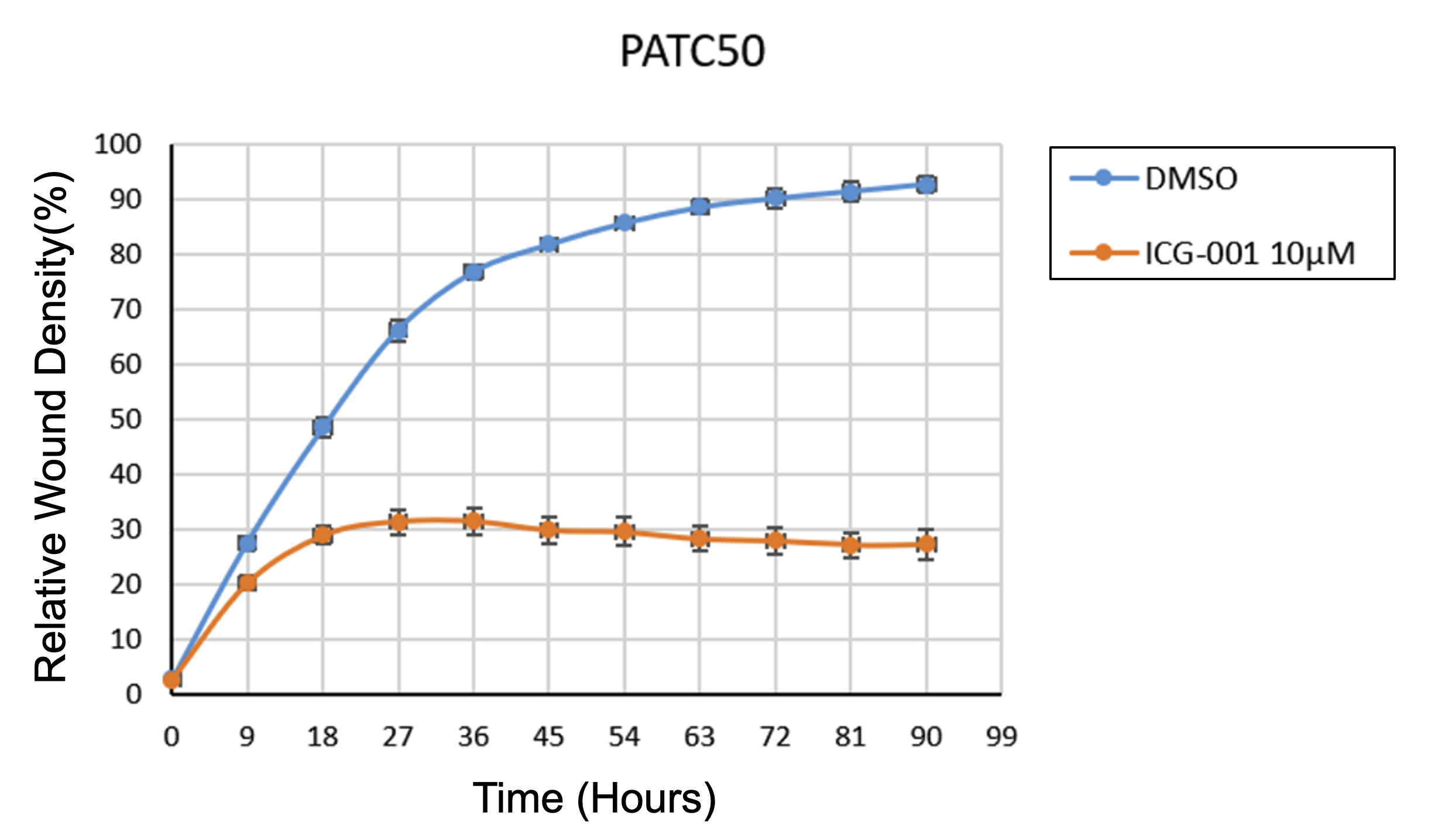

**B**

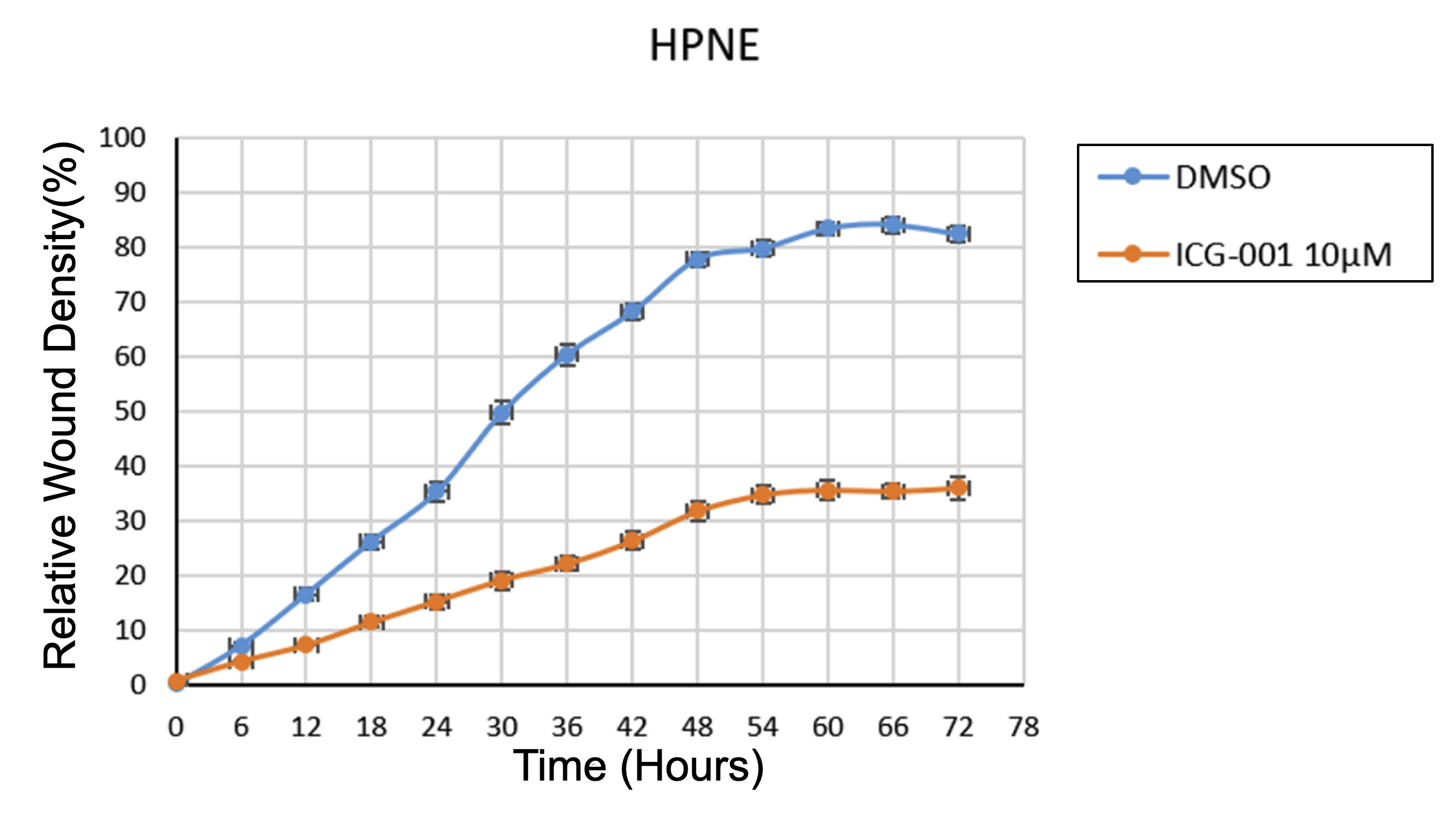

**Supplementary Figure S3. Apoptosis analysis of cells with the treatment of ICG-001.**

(**A**) Apoptosis analysis of PATC50 cells with ICG-001 treatment. (**B**) Apoptosis analysis of HPNE cells with ICG-001 treatment. * p < 0.05, student t test.

**A**

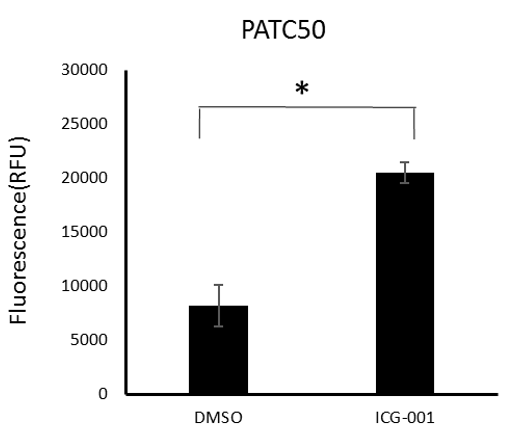

**B**

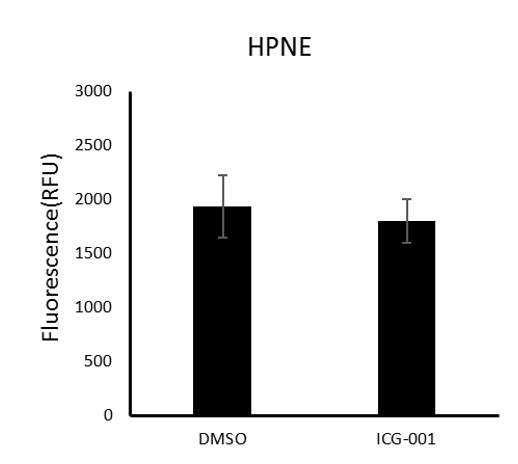

**Supplementary Figure S4. Cell growth status with the combined treatment.**

(**A**) Combination treatments of ICG-001 and Gemcitabine in PANC1 and PATC53. (**B**) Growth status of PATC50 cells with the combined treatment of ICG-001 and Gemcitabine. (**C**) Growth status of HPNE cells with the combined treatment of ICG-001 and Gemcitabine. I: ICG-001, G: Gemcitabine.

**A**

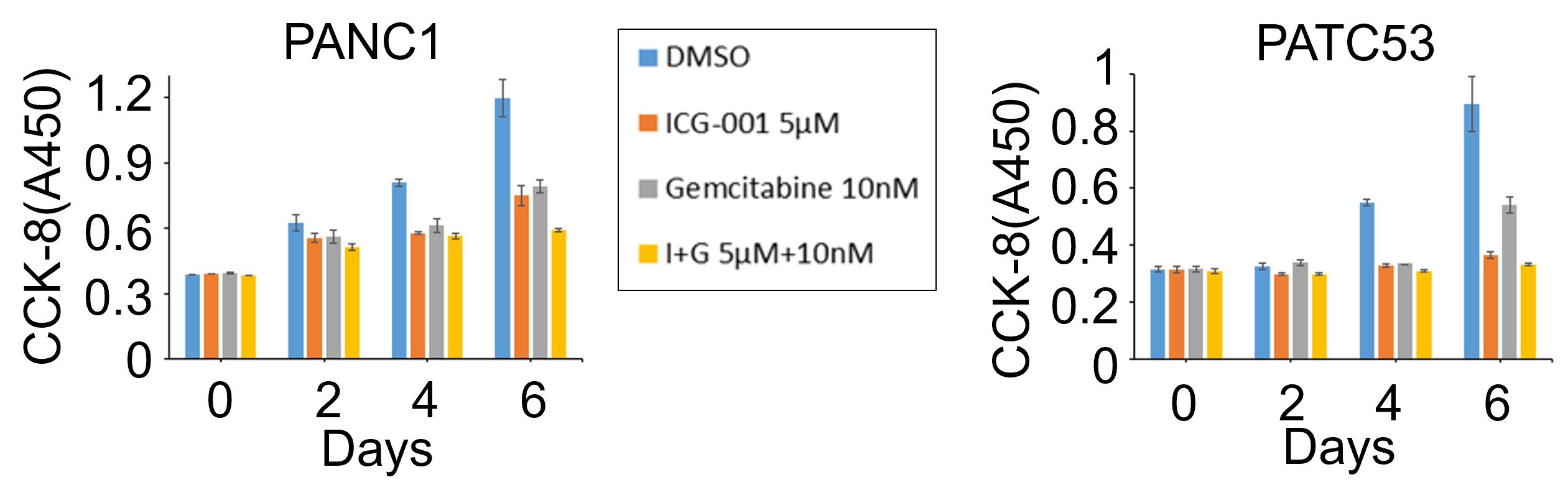

**B**

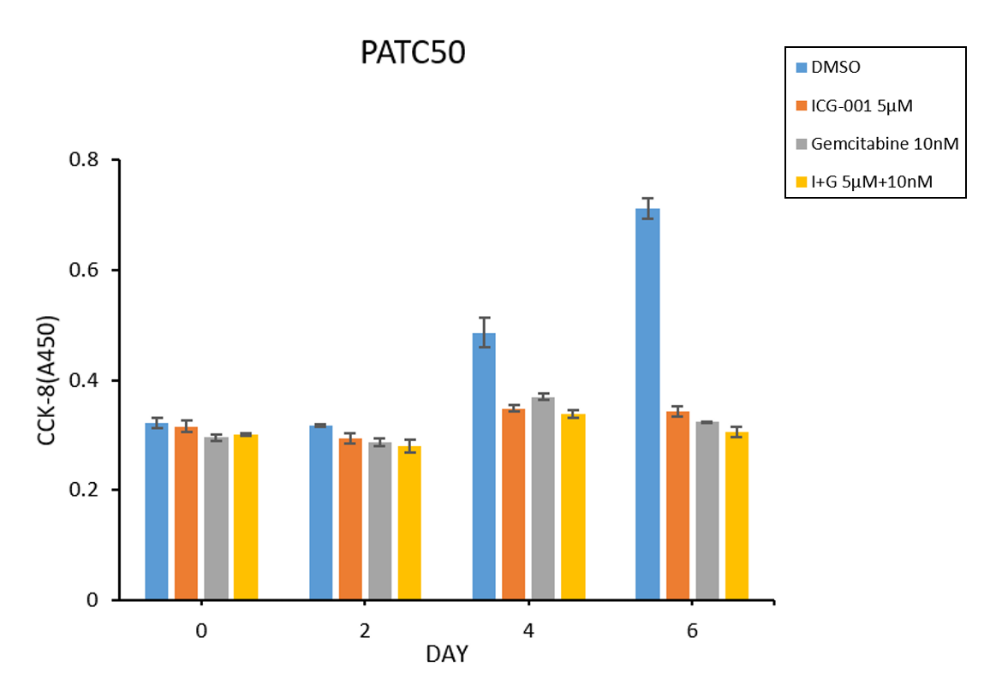

**C**

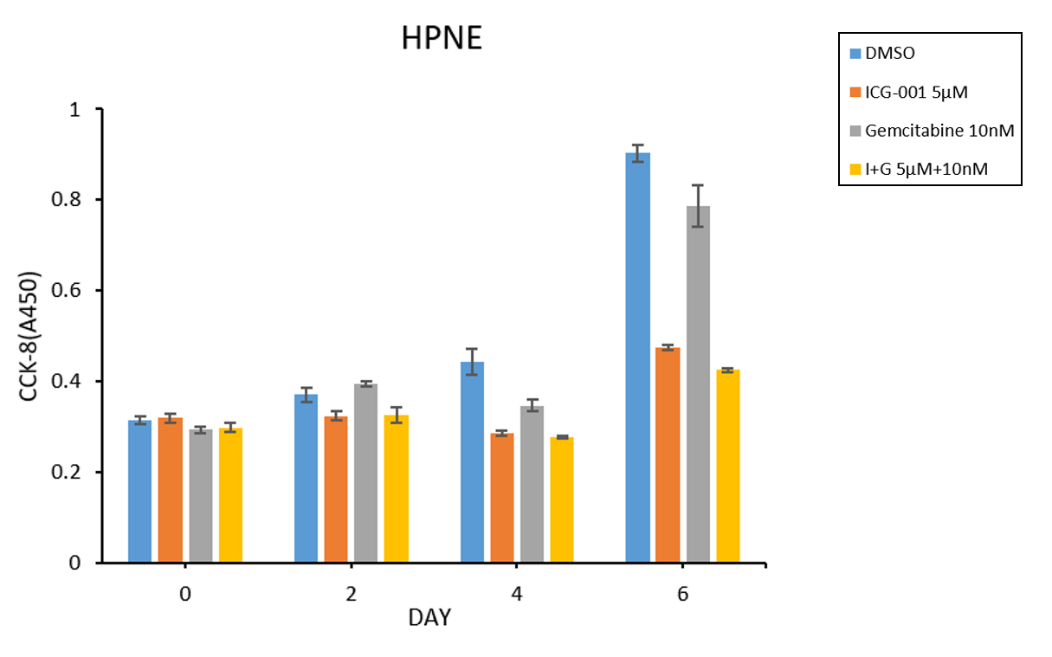

**Supplementary Figure S5. The repeated Western blotting in PATC53 cells with the deletion of the enhancer region of IRS1.**

(**A**) Three technical replicates of Western blotting were performed to show the expression levels of the downstream proteins in insulin signaling pathway. (**B**) No significant difference was showed by paired student t-test when Del-01 and Del-02 were treated with ICG-001 comparing to sgEmpty control.

**A**

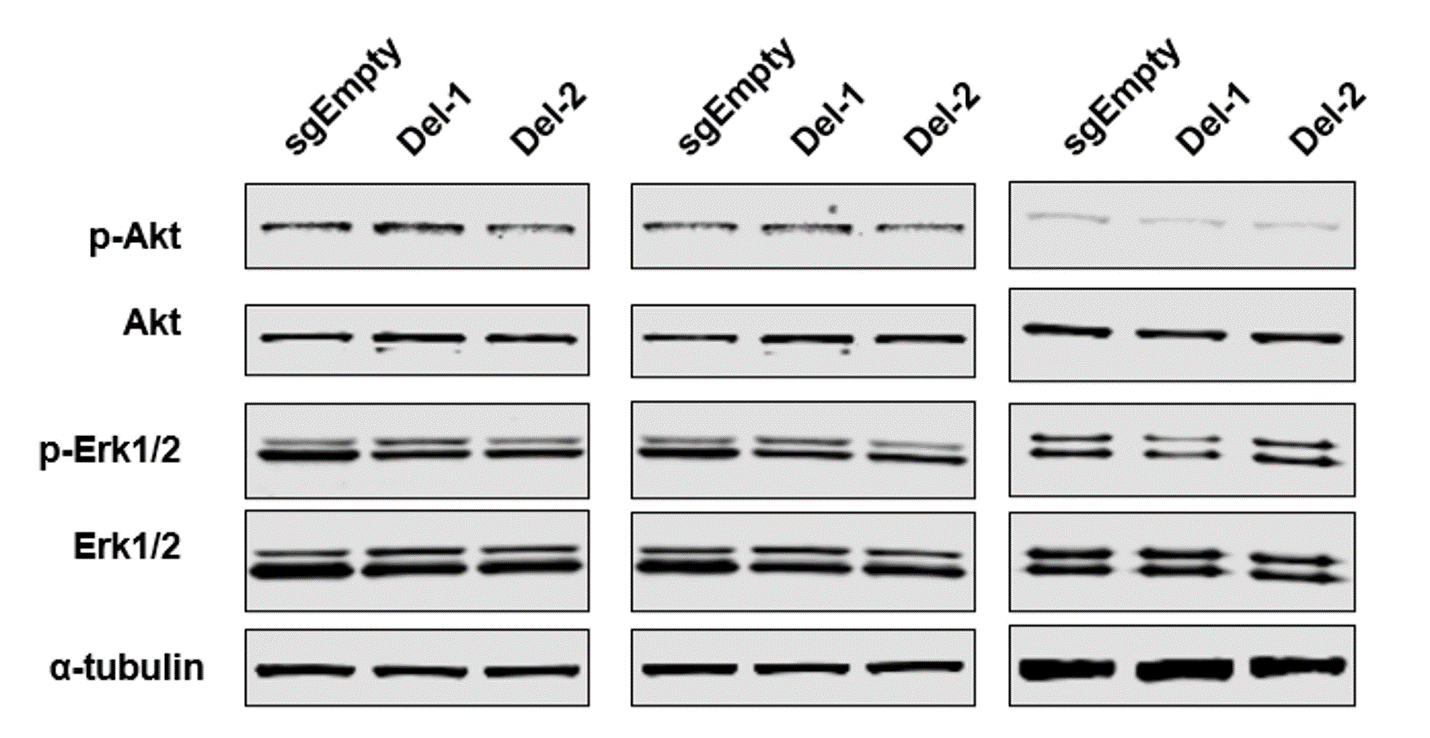

**B**

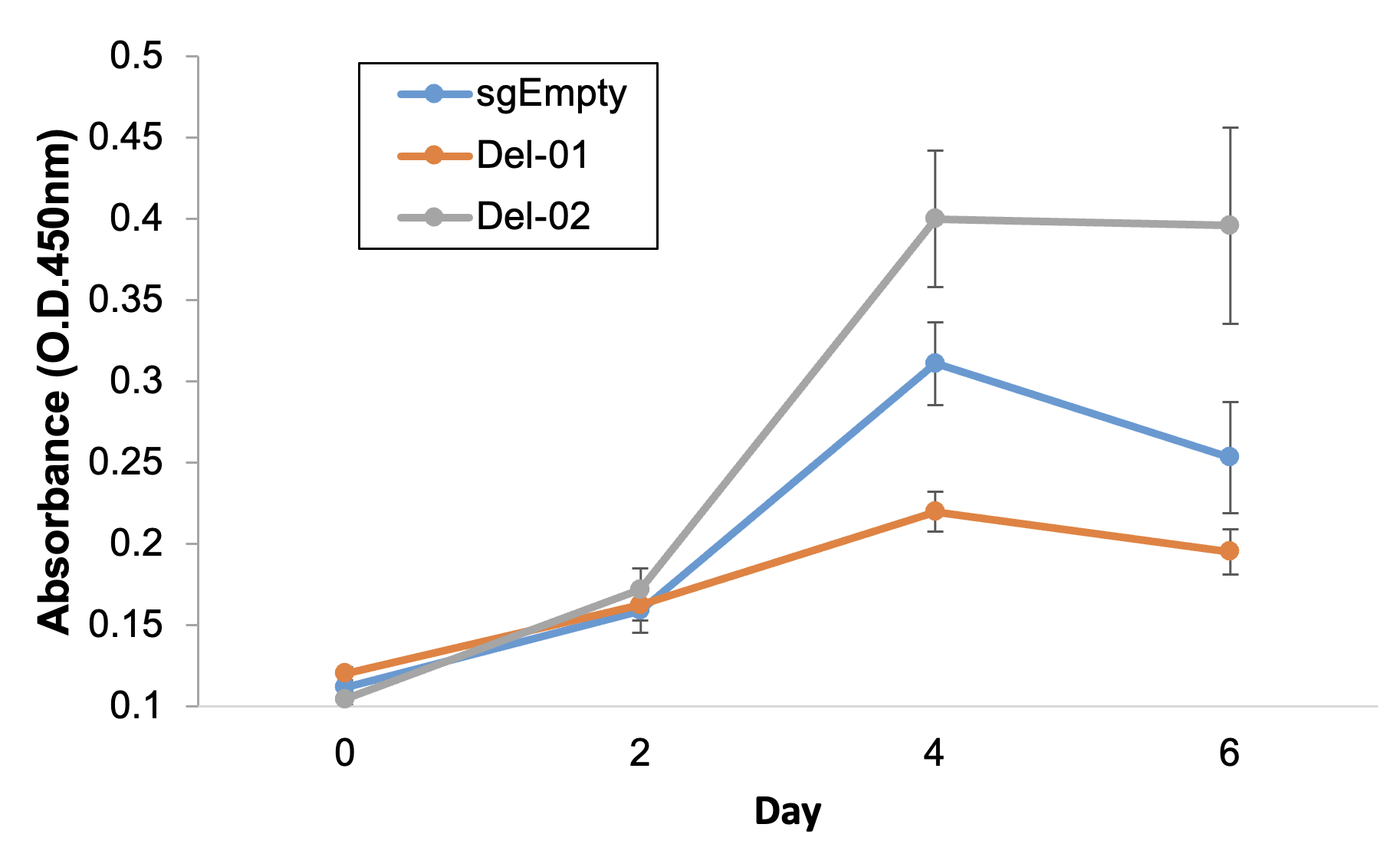

**Supplementary Figure S6. RHBDD1 does not influence the cell growth and the mRNA level of IRS1 in PATC53 cells.**

(**A**) RT-qPCR showed the mRNA expression level of RHBDD1 of the enhancer deletion clones (Del-01 and Del-02) decreased significantly. The deleted enhancer region of IRS1 was located in the coding region of the gene RHBDD1. (**B**) To check if the decreased level of RHBDD1 imfluences the the cell growth in PATC53 cells, shRNAs (shControl or shRHBDD1) were used to deplete the mRNA level of RHBDD1. (**C**) No significant differece of the cell growth curves was observed between PATC53 cells without or with RHBDD1 depletion using shRNA (shControl or shRHBDD1). (**D**) The mRNA level of IRS1 was not inhibited or increased in PATC53 cells with RHBDD1 depletion.

**A**

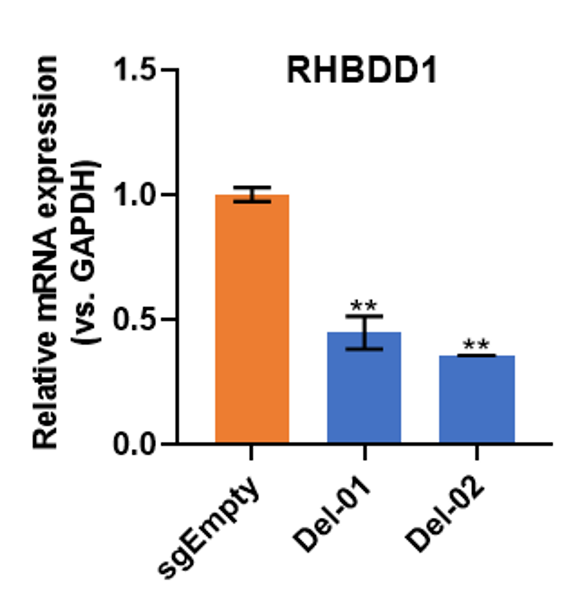

**B**

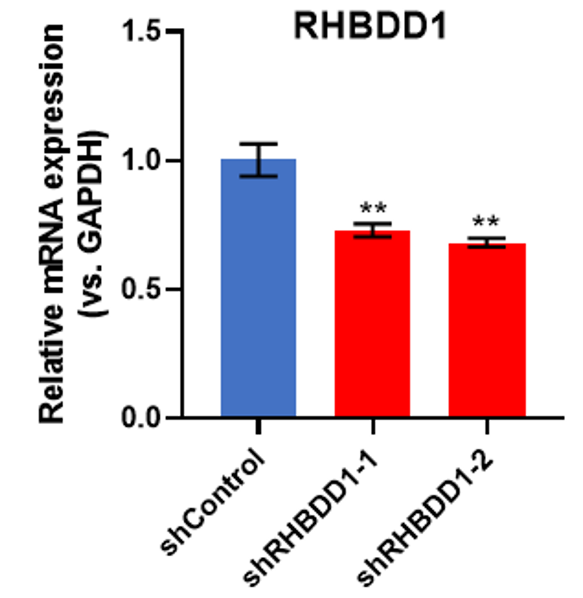

**C**

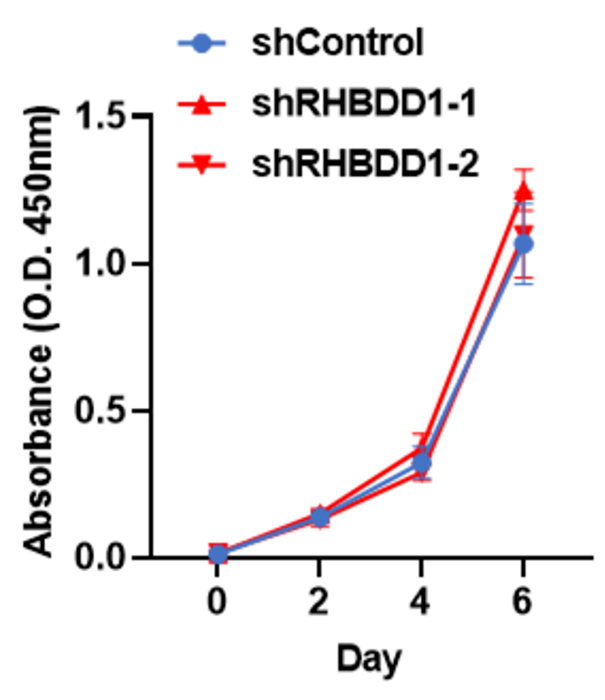

**D**

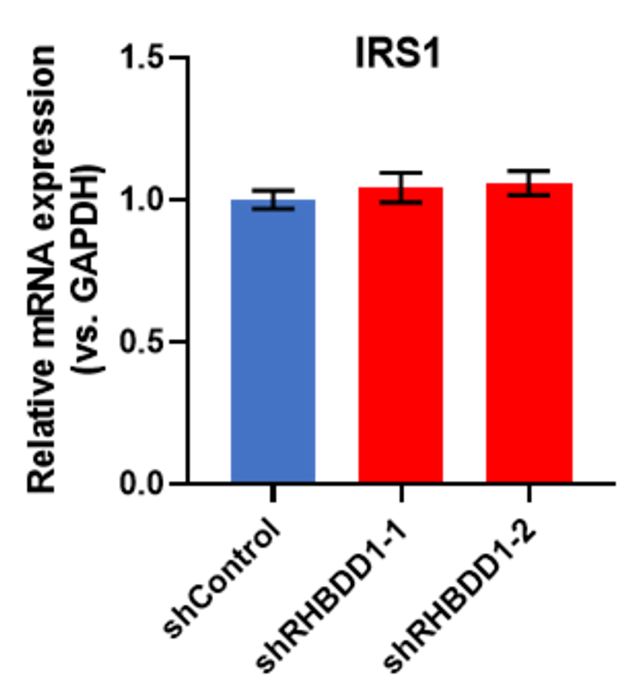

**Supplementary Table S1. Reads of Hi-C data**

PANC1

|  | ENCODE PANC1 | ENCODE PANC1 | PANC1+ICG-001 | PANC1+ICG-001 |
| --- | --- | --- | --- | --- |
|  | Rep1 | Rep2 | Rep1 | Rep2 |
| Raw reads | 151,103,112 | 137,874,940 | 138,855,403 | 160,655,787 |
| Uniquely mapped | 90,313,153 | 67,559,617 | 84,100,007 | 96,896,977 |
|  | (59.80%) | (49.00%) | (60.60%) | (60.30%) |
| Valid Pairs | 82,484,838 | 62,661,214 | 73,572,169 | 87,263,424 |
|  | (91.30%) | (92.70%) | (87.50%) | (90.10%) |
| Removing duplicates | 81,397,598 | 56,817,809 | 67,848,364 | 79,953,474 |
|  | (98.68%) | (90.67%) | (92.22%) | (91.62%) |

PATC53

|  | PATC53 | PATC53 | PATC53+ICG-001 | PATC53+ICG-001 |
| --- | --- | --- | --- | --- |
|  | Rep1 | Rep2 | Rep1 | Rep2 |
| Raw reads | 335,546,131 | 334,793,995 | 338,795,426 | 284,332,904 |
| Uniquely mapped | 189,132,486 | 178,395,348 | 192,697,078 | 136,147,287 |
|  | (56.40%) | (53.30%) | (56.90%) | (47.90%) |
| Valid Pairs | 111,736,560 | 69,126,040 | 123,780,415 | 74,636,388 |
|  | (59.10%) | (38.70%) | (64.20%) | (54.80%) |
| Remove duplicates | 101,494,245 | 62,896,779 | 112,875,760 | 71,342,891 |
|  | (90.83%) | (90.99%) | (91.19%) | (95.59%) |

**Supplementary Table S2. Reads of RNA-seq data**

PANC1

|  | PANC1 Rep1 | PANC1 Rep2 | PANC1 Rep3 | PANC1_ICG-001 Rep1 | PANC1_ICG-001 Rep2 | PANC1_ICG-001 Rep3 |
| --- | --- | --- | --- | --- | --- | --- |
| Raw Reads | 33,744,666 | 46,629,164 | 37,984,529 | 38,716,394 | 45,844,799 | 38,864,612 |
| Uniquely Aligned | 28,340,835 (83.99%) | 37,937,998 (81.36%) | 30,678,396 (80.77%) | 33,270,022 (85.93%) | 36,975,473 (80.65%) | 32,398,583 (83.36%) |

PATC53

|  | PATC53 Rep1 | PATC53 Rep2 | PATC53 Rep3 | PATC53_ICG-001 Rep1 | PATC53_ICG-001 Rep2 | PATC53_ICG-001 Rep3 |
| --- | --- | --- | --- | --- | --- | --- |
| Raw Reads | 47,482,728 | 47,497,802 | 35,666,123 | 48,120,131 | 47,915,491 | 44,310,149 |
| Uniquely Aligned | 39,734,767 (83.68%) | 38,895,089 (81.89%) | 29,056,894 (81.47%) | 40,314,629 (83.78%) | 38,605,972 (80.57%) | 35,973,406 (81.19%) |

HPNE

|  | HPNE Rep1 | HPNE Rep2 | HPNE Rep3 | HPNE_ICG-001 Rep1 | HPNE_ICG-001 Rep2 | HPNE_ICG-001 Rep3 |
| --- | --- | --- | --- | --- | --- | --- |
| Raw Reads | 34,342,760 | 37,379,750 | 38,072,940 | 39,377,087 | 43,773,283 | 41,386,362 |
| Uniquely Aligned | 29,508,224 (85.92%) | 31,116,071 (83.24%) | 29,528,802 (77.56%) | 32,458,878 (82.43%) | 36,792,886 (84.05%) | 33,944,167 (82.02%) |

PATC50

|  | PATC50 Rep1 | PATC50 Rep2 | PATC50 Rep3 | PATC50_ICG-001 Rep1 | PATC50_ICG-001 Rep2 | PATC50_ICG-001 Rep3 |
| --- | --- | --- | --- | --- | --- | --- |
| Raw Reads | 46,551,843 | 37,445,665 | 40,955,126 | 46,845,888 | 44,896,703 | 35,538,038 |
| Uniquely Aligned | 39,037,119 (83.86%) | 29,917,096 (79.89%) | 31,082,866 (75.89%) | 39,712,849 (84.77%) | 36,694,714 (81.73%) | 28,958,654 (81.49%) |

**Supplementary Table S3. List of sgRNA sequences and RT-qPCR primers**

| **Oligonucleotides** | **Sequence** | **Experiment** |
| --- | --- | --- |
| IRS1 disDel-sgRNA-F-oligo1 | caccgTGACCATGGAAGTTCTACCC | CRISPR-KO sgRNA design |
| IRS1 disDel-sgRNA-F-oligo2 | aaacGGGTAGAACTTCCATGGTCAc | CRISPR-KO sgRNA design |
| IRS1 disDel-sgRNA-R-oligo1 | caccgAGTAATTGCACTATACTACC | CRISPR-KO sgRNA design |
| IRS1 disDel-sgRNA-R-oligo2 | aaacGGTAGTATAGTGCAATTACTc | CRISPR-KO sgRNA design |
| IRS1 disDel Validation-in-F | TTGCTGCTTTTCCTGTAAACCTTAG | CRISPR-KO validation primer |
| IRS1 disDel Validation-in-R | AAGCTCAACTGACAATGAGAAACTG | CRISPR-KO validation primer |
| IRS1 disDel Validation-out-F | TGGAATGCTGTTCCATTTAACTCAG | CRISPR-KO validation primer and Sanger seq |
| IRS1 disDel Validation-out-R | CCTTGGGGAGCTAGTGAAAACATA | CRISPR-KO validation primer |
| IRS1-Forward | ACTGGACATCACAGCAGAATGA | RT-qPCR |
| IRS1-Reverse | TCGTACCATCTACTGATGAGGAAG | RT-qPCR |
| RHBDD1-Forward | GTGCTGTAGGTTTCTCAGGAGTT | RT-qPCR |
| RHBDD1-Reverse | ACAGGAAAGCCCAAAATGTTGAC | RT-qPCR |

**Supplementary Table S4. Plasmids used in this study**

| **Plasmid Name** | **Company** | **Catalog** | **Experiments** |
| --- | --- | --- | --- |
| lentiCRISPR v2 plasmid | Addgene | #52961 | CRISPR/Cas9 mediated distal region deletion |
| RHBDD1 shRNA Plasmid (h) | Santa Cruz Biotechnology | sc-94654-SH | shRNA |
| Control shRNA Plasmid-A | Santa Cruz Biotechnology | sc-108060 | shRNA |

**Supplementary Table S5. List of 3C-qPCR primers**

| **No** | **Name** | **Primer** |
| --- | --- | --- |
| 1 | EIF4EBP1 _Anchor | TATTTGAGTCTGTGGAGTCTTAAAAATATC |
| 2 | EIF4EBP1 | GTATTAGGTTTCATTATACACATTTGAGAG |
| 3 | FBP1_Anchor | GACTCTGTCTGAAAAATAAAGAAATATAGC |
| 4 | FBP1 | ATATATTTATGCTAGTTAGAGTTACGGTTC |
| 5 | FBP2_Anchor | CAAGTACAAGTCATGATAACACAATACATA |
| 6 | FBP2 | ACATAAGGGAGAGTATAAGGGAGAGTATTA |
| 7 | IRS1_Anchor | CTTTTATCATACATGACAGTACCTTATACT |
| 8 | IRS1 | AACTTCTAATATACAAGCATCTACTTACAC |
| 9 | KRAS_Anchor | TAGTAGATTAGTACACCACGTAGTATCTTT |
| 10 | KRAS | GTGTTTACAGTATTATTTAAGAAGCCTATG |
| 11 | MAPK9_Anchor | CAACTGATCTGTAACTAAGAATTCTACACA |
| 12 | MAPK9 | CTGTTTATACATAAAGATACTTCTGAATGG |
| 13 | MKNK1_Anchor | CGTTCAGTAGACTTAAGATGGTTTAATAAT |
| 14 | MKNK1 | GTATAACCACCACTATAATCAGAAATACAG |
| 15 | NRAS_Anchor | TTACTTTCTCTCCTCTTATTCCTTTAATAC |
| 16 | NRAS | CTTACTGAGAACTTAATTTTGAATGTCTAC |
| 17 | PIK3R1_Anchor | GTAAGTAGAGAGTGATGGTTATAACATTTG |
| 18 | PIK3R1 | CTTGTATCTTGTAGACAAACTTGTAAAAAC |
| 19 | PPP1CB_Anchor | CTTAGAACTAAAAACCAAAGACTTCTACTT |
| 20 | PPP1CB | TCACAGTACTATTTTATTCAAGTGTCTAAC |
| 21 | PPP1R3A_Anchor | CTATTCATAGGCATAAAGAGATAATAAGGT |
| 22 | PPP1R3A | CTAAGTAAGTTATACGACATATTCATTTCC |
| 23 | PRKAA2_Anchor | CTGATGTTTAACTATGTGATCTATTTTGAG |
| 24 | PRKAA2 | CCTACTACTTGAGTAAGTTATTTCACATCT |
| 25 | PRKAR1A_Anchor | GAATTATAGTAGATATTCTGGCTTTCTCTT |
| 26 | PRKAR1A | GCTAAAAGAGTCTCTCTCTAAGTATGTAAT |
| 27 | PRKAR1B_Anchor | ACTAGAAGTCAGTAACAGAAAGATAGATAG |
| 28 | PRKAR1B | CGATACTTGTAGGTGTAGTACAATTTTTAT |
| 29 | PRKCZ_Anchor | TATAGCAATGTGTAAAGAAGTGTGATTAAC |
| 30 | PRKCZ | AATACTGATGTTGGAAACTAGAAAACTATG |
| 31 | PTPN1_Anchor | AAAGTTTATAGGAGCTTTGTGAGTATAGTT |
| 32 | PTPN1 | TACGTTTAGGTATGTTTAGAGACAGATACT |
| 33 | PYGM_Anchor | CTTCAGTTCTATATTACTTCTTTCTTTTCC |
| 34 | PYGM | CTTACTTTGATTGTATCATTAGAGTTGATG |
| 35 | RAF1_Anchor | AATTCAACTAGTTCACTATCTACAACAAAG |
| 36 | RAF1 | CACTACACACTTATTCTAATAGCTAAAATC |
| 37 | GAPDH_Anchor | ATGCAAGGCTTTCTCTTAAATTAGC |
| 38 | GAPDH | AATTCTGAGCATTCTGTAGCAAACT |

**Supplementary Table S6. Antibodies used for Western blotting experiments**

| **Antibody Name** | **Company** | **Catalog** | **Dilution** |
| --- | --- | --- | --- |
| Rabbit monoclonal anti-IRS1 | Abcam | ab40777 | 1:2000 |
| Rabbit monoclonal anti-β-actin | Abcam | ab227387 | 1:5000 |
| Mouse monoclonal anti-α-tubulin | Cell Signaling Technology | 3873 | 1:2000 |
| Mouse monoclonal anti-Akt (pan) | Cell Signaling Technology | 2920 | 1:2000 |
| Rabbit polyclonal anti-Phospho-Akt (Ser473) | Cell Signaling Technology | 9271 | 1:2000 |
| Mouse monoclonal anti-p44/42 MAPK (Erk1/2) | Cell Signaling Technology | 4696 | 1:2000 |
| Rabbit monoclonal anti-phospho-p44/42 MAPK (Erk1/2) (Thr202/Tyr204) | Cell Signaling Technology | 4370 | 1:2000 |
